# Fibroblast response to burn injury in larval zebrafish mirrors developmental maturation and is inhibited by infiltrating neutrophils

**DOI:** 10.64898/2026.06.19.733402

**Authors:** Adam Horn, Yiran Hou, Jayne M. Squirrell, Alexandra M. Fister, Julie Rindy, Veronika Miskolci, Jonathan H. Schrope, Russell Burke, Colin Dewey, Kevin Eliceiri, Anna Huttenlocher

## Abstract

Fibroblasts mediate tissue repair after damage, but aberrant fibroblast behavior in response to injury can result in impaired wound healing. Severe burn injury often results in tissue scarring, but the underlying mechanisms by which fibroblasts respond to burn injury and the role of inflammation in fibrosis are not well understood. Here we developed fluorescent reporters of collagen expressing mesenchymal cells enabling real time imaging of fibroblasts during homeostatic development and in response to burn injury using larval zebrafish. We find that fibroblasts derived from the mesenchyme respond to burn injury by engaging in a maturation process, characterized by the expression of vimentin, which is reminiscent of larval development. In burned tissue, fibroblast maturation is perturbed by prolonged neutrophil infiltration, resulting in disorganized extracellular matrix (ECM) and delayed ECM remodeling, which can be rescued by neutrophil depletion. This work adds to our understanding of fibroblast development in zebrafish and shows that collagen expressing mesenchymal cells regulate ECM remodeling in coordination with immune cells during burn wound healing.

**Summary Statement:** Vimentin-positive fibroblasts are required for normal wound healing in larval zebrafish, but the presence of inflammatory neutrophils after burn injury impairs fibroblast maturation and collagen remodeling.

## Introduction

Fibroblasts mediate tissue organization and remodeling during development and wound healing [1]. In response to tissue damage, fibroblasts are activated, enabling their migration to the site of injury [2, 3]. During the remodeling phase of tissue repair, fibroblasts produce extracellular matrix (ECM), including collagen, which is required for normal wound healing [4]. Importantly, fibroblasts simultaneously engage in ECM remodeling through the secretion of ECM degrading enzymes, such as matrix metalloproteinases [1]. Precise regulation of ECM deposition and remodeling is required for normal healing and to prevent the buildup of scar tissue. While much is known about the function of fibroblasts *in vitro*, dissecting their role *in vivo* has proved challenging due to the functional heterogeneity fibroblasts display within tissues [5–8].

Larval zebrafish have proven to be a useful model organism to investigate the developmental heterogeneity of fibroblasts and their response to tissue damage. Recent advances in characterizing fibroblast development using zebrafish have identified that mesenchyme-derived cells of the developing fin fold display conserved molecular and functional characteristics of mammalian fibroblasts [9–12], and contribute to early damage signaling in wounded tissue [13]. Despite these studies, there is a need to generate additional reporters of mesenchyme-derived fibroblasts to better understand their function in response to tissue damage *in vivo*.

Burn wounding often results in scarring and subsequently non-functional tissue [14, 15], but the underlying mechanisms of how fibroblasts respond to injury and contribute to scar formation in burned tissue are not well understood. We have previously used the larval zebrafish as a model to understand the pathophysiology of burn wound healing. Burn wounding of larval zebrafish recapitulates the hallmarks of burn wound pathology including a chronic and excessive inflammatory response, impaired sensory function, and susceptibility to infection [16–19]. Additionally, we previously identified a subpopulation of mesenchymal cells expressing the fibroblast marker vimentin that show an impaired response immediately following burn injury [20], but the behavior of these cells during burn wound healing remains largely unknown.

Fibroblasts undergo a cell-intrinsic activation process, mediated in part by increased expression of the intermediate filament protein vimentin [21–23], which in turn promotes cell migration and morphological changes associated with altered fibroblast function [24–27]. Vimentin expression is required for injury-induced expression of collagen [20, 24] and regulates normal wound healing in both burn and mechanical wounds [24, 28, 29]. In addition to cell-intrinsic cues, the surrounding tissue environment also modulates fibroblast activity during wound healing [2, 3]. In particular, the inflammatory response has been shown to play an important role in determining the severity of tissue scarring after injury [30]. In normal wound healing, the inflammatory phase of tissue repair is transient, with inflammation resolving prior to the onset of tissue remodeling. However, after burn wound injury prolonged inflammation can overlap temporally with ECM deposition and remodeling. Burn wounding is known to produce a particularly extensive neutrophil response, in which neutrophils are thought to contribute to progressive tissue damage [31]. Further, neutrophils are more easily able to infiltrate burned tissue than other innate immune cells and fail to resolve efficiently compared to injury caused by mechanical trauma [32, 33]. While excessive neutrophil inflammation is associated with fibrosis, neutrophils also positively regulate fibroblasts, such that neutrophil depletion has also been shown to contribute to fibrosis [34, 35]. Thus, additional work is required to elucidate the precise interplay between infiltrating neutrophils and fibroblasts within the wound microenvironment.

Here, we used unbiased RNA-sequencing and live imaging techniques to investigate how fibroblast behavior contributes to burn wound healing. RNA-sequencing of burned larvae identified temporally overlapping molecular signatures related to the innate immune response and fibroblast activity in burn wounded tissue throughout the healing process. To directly visualize ECM producing cells, we generated a transgenic reporter zebrafish to label cells expressing collagen type 1 alpha 1a (col1a1a), an abundant collagen comprising actinotrichia in the developing caudal fin [36]. Simultaneous imaging of col1a1a-expressing cells with our previously validated vimentin reporter zebrafish revealed a distinct subpopulation of fibroblasts in the developing mesenchyme that respond to burn injury. These cells are modulated by the inflammatory response to burn, such that the presence of neutrophils impairs ECM production and remodeling. This work improves our understanding of larval zebrafish mesenchymal cell dynamics during development and tissue repair and shows that fibroblast interactions with infiltrating neutrophils are inhibitory to burn wound healing.

## Results

### RNA-sequencing identifies genes associated with fibroblast identity and prolonged immune response in burned tissue

To probe for pathways that may alter fibroblast function and modify ECM remodeling during burn wound healing, we performed bulk RNA-sequencing analysis in larval zebrafish. We burned the caudal fins of 3 day-post-fertilization (dpf) larvae and collected wounded tissue at 6-, 24-, 48-, and 72-hours post-burn (hpb), together with unwounded, age-matched fins as controls (Fig. 1A). We first identified the presence of differentially expressed genes at each time point, which were chosen to represent the full spectrum from early tissue damage responses (6 hpb) to late-stage healing (72 hpb) (Fig. 1B-E, Supplemental Table 1). Notably, we found the upregulation of genes related to fibroblast function, including *ecm2*, *mmp11b*, *mmp13b*, and *htra1b*, at each time point during burn wound healing. This confirmed a canonical fibroblast response is present following burn injury in larval zebrafish.

**Figure 1:**
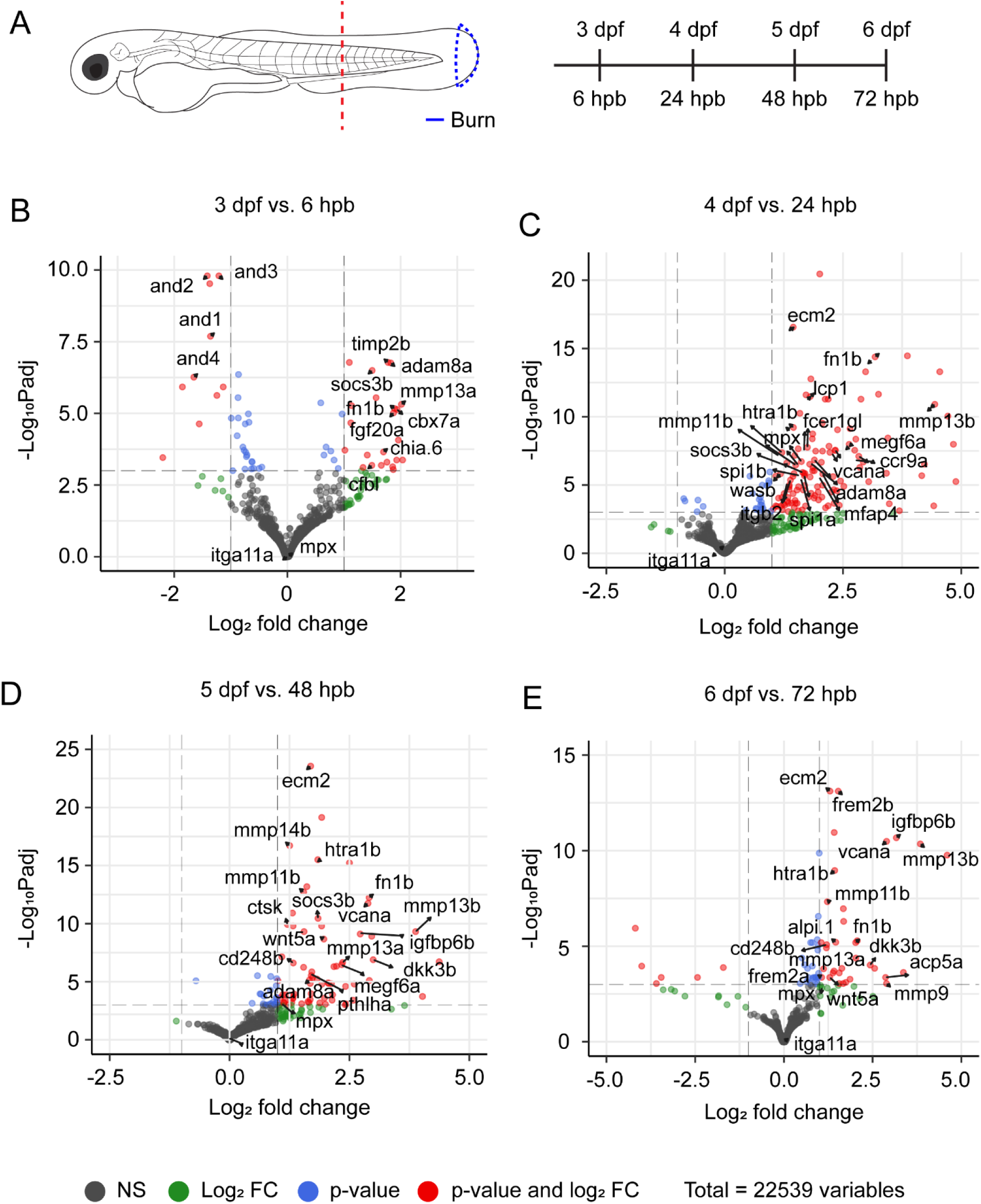
Gene expression profile of burn wound healing. (A) Schematic of sample collection for bulk RNA-sequencing. Zebrafish larvae were burnt at the tip of the caudal fin (blue dotted region) at 3 dpf. Tail fin samples containing the burn wound until the red dotted line were collected at 3/4/5/6 dpf and 6/24/48/72 hpb. (B-E) Volcano plots showing differentially expressed genes comparing between burnt and unwounded controls at 3 dpf/6 hpb (B) 4 dpf/24 hpb (C) 5 dpf/48 hpb (D) and 6 dpf/72 hpb (E). Genes related to fibroblast and myeloid cell identity are highlighted. Adjusted p-value cutoff is set at 1e-3 for visualization purposes, while threshold for log2 fold change is set at 1.

To identify potential pathways that modify ECM remodeling during burn wound healing, we next performed Weighted Gene Co-Expression Network Analysis (WGCNA) across all samples (Supplemental Figure 1A) [37]. We calculated eigengene expression changes by aligning gene modules with metadata, allowing us to identify modules containing genes that show positive expression changes in burn wounding (Modules-5/-8/-14). Eigengenes are modules of co-expressed genes. We were able to infer the functions of each gene module through Gene Ontology (GO) enrichment analysis (Supplemental Table 2) [38]. Module-8 illustrates functions closely related to ECM remodeling (GO:0060348 bone development, GO:0030198 extracellular matrix organization, and GO:0048729 tissue morphogenesis), corroborating the presence of a fibroblast response related to tissue repair over time. Interestingly, the other two modules showed sustained enrichment for the regulation of the immune response, with Module-14 limited to myeloid leukocyte differentiation (GO:0002573) and Module-5 being associated with a wider range of immune functions (GO:0002682 regulation of immune system process, GO:0050900 leukocyte migration, and GO:0006954 inflammatory response). These data suggested the overlapping presence of both fibroblast and innate immune activity over time following burn injury. Based on this, we hypothesized that innate immune interactions with fibroblasts may modify their wound-induced function and subsequent ECM remodeling in burned tissue.

### Mesenchymal cells express col1a1a and vimentin during zebrafish development

To begin understanding the role of fibroblasts and their regulation in response to burn injury, we first generated a transgenic reporter zebrafish – *Tg(Col1a1a:zmCherry)* – driving fluorescent protein expression under the control of the *col1a1a* promoter, which is a marker of fibroblast identity and an abundant collagen protein comprising actinotrichia in the developing caudal tailfin [36, 39]. Live imaging of *Tg(Col1a1a:zmCherry)* larvae identified notable populations of mesenchymal cells during development, specifically in the region surrounding the eye, pectoral fin, trunk, and tail fin (Fig. 2A). While col1a1a-positive cells are present throughout the tail fin, we noted a distinct difference in morphology based on their position extending away from the midline. Cells toward the interior appeared flat and circular, while cells extending outward toward the fin edge took on the intricate, elongated morphology previously associated with fibroblasts of the tail fin mesenchyme [9, 13, 40].

**Figure 2:**
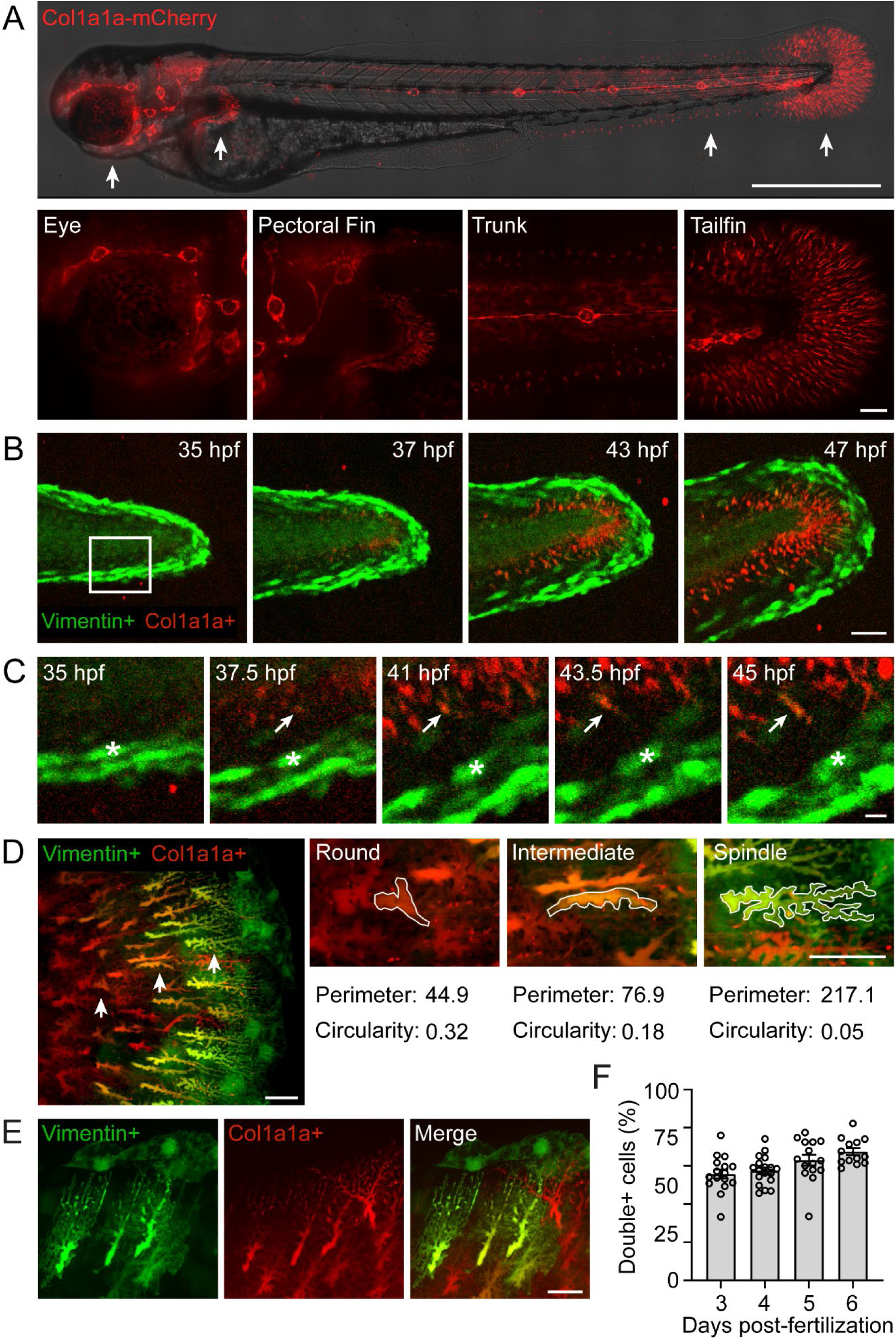
Vimentin expression is associated with fibroblast maturation during development. (A) Image showing *Tg(Col1a1a:zmCherry)* larvae 3 days post-fertilization. Arrows point to regions of high *col1a1a* expression shown in insets below. Scale bar = 500 µm (top) and 20 µm (insets). (B) Time series showing *Tg(Col1a1a:zmCherry)xTg(−2vim:eGFP)* double-positive larva at the indicated number of hours post-fertilization (hpf). White box indicates area shown in D. Scale bar = 50 µm. (C) Time series showing cropped region of larva from C. Asterisk indicates a single vimentin-positive, col1a1a-negative cell. Arrow indicates the same col1a1a positive cell over time. Scale bar = 10 µm. (D) Image showing *Tg(Col1a1a:zmCherry)xTg(−2vim:eGFP)* double-positive larvae 3 days post-fertilization. Arrows point to cells depicted in inset at right. Perimeter and circularity are reported for individual traced cells. Scale bar = 20 µm. (E) High magnification image of *Tg(Col1a1a:zmCherry)xTg(−2vim:eGFP)* double-positive larva 3 days post-fertilization. Image is taken in the tail fin region. Scale bar = 20 µm. (F) Quantification showing the percentage of tail fin fibroblasts that are double positive for col1a1a and vimentin at the indicated day post-fertilization. N ≥ 16 larvae for each day post-fertilization.

In addition to our investigation of col1a1a, we also developed transgenic reporter fish lines for cells expressing other type I collagen – collagen 1a1b and collagen 1a2 (Supplemental Figure 2A-D). Each of these labeled mesenchymal cells of the larval zebrafish tail fin, which we confirmed by identifying the localization of collagen-expressing cells to be sub-epithelial (Supplemental Fig 2E). Further investigation of col1a2 expressing cells revealed loss of expression in the tail fin during larval development, while there was complete overlap in expression of col1a1b and col1a1a in cells localized to the tail fin (Supplemental Fig. 1F). For these reasons, we continued our investigation of fibroblasts by focusing on the col1a1a reporter line.

We previously found that vimentin expressing mesenchymal cells respond to tissue injury and that vimentin expression is required for increased collagen expression in wounded tissue [20], but whether these cells were related to collagen expressing fibroblasts remained unclear. Crossing *Tg(Col1a1a:zmCherry)* and *Tg(−2vim:eGFP)* allowed us to assess the developmental relationship between vimentin-expression and col1a1a-positive mesenchymal cells. Taking advantage of a distinct population of vimentin-positive, but col1a1a-negative, cells lining the outer boundary of the tailfin, we performed live imaging of larval development starting on the first day post-fertilization. At this stage, no col1a1a-positive mesenchymal cells are present, but vimentin-positive cells already lined the outer boundary of the tailfin allowing for tracking of individual cells over time (Fig. 2B, C, Supplemental Fig. 2G). Starting roughly 37.5 hours post-fertilization, fluorescence indicating the expression of col1a1a began to appear, notably in cells that were not vimentin positive (Fig. 2C, Supplemental Fig. 2G). These cells initially expressed no vimentin and only began expressing vimentin hours later.

By 3 days post-fertilization, we observed what appeared to be developmentally mature fibroblasts lining the outer fin edge, consistent with observations from other groups [13]. Intriguingly, there were clear sub-populations of mesenchymal cells based on their position relative to the midline. While we found no vimentin expression in col1a1a-positive cells closest to the midline of the developing larvae, a clear and progressive increase in vimentin expression was observed in cells extending outward toward the edge of the fin (Fig. 2D). This also corresponded with a clear change in morphology, which is consistent with a role for increased vimentin expression in regulating fibroblast morphology [21, 41]. Indeed, we found that increased vimentin-expression in cells closer to the fin edge was associated with increased perimeter and decreased circularity (Fig. 2D).

Finally, we sought to quantify the proportion of col1a1a- and vimentin-double positive fibroblasts over the course of early tail fin development (Fig. 2E). This revealed a progressive increase in the percentage of double positive fibroblasts over time, with 55.7±2.4% of fibroblasts expressing both markers at 3 days post-fertilization increasing to 67.4±1.9 by 6 days post-fertilization (Fig. 2F). Notably, this increase in the proportion of double positive fibroblasts was driven primarily by an increase in vimentin expression, as the number of total col1a1a-positive fibroblasts did not increase over time, despite a clear growth in tail fin size (Supplemental Fig. 2H-I). Closer examination of the tail fin boundary revealed the persistence of vimentin-positive, but col1a1a-negative cells (Fig. 2E). These cells, which were present as early as 1 day post-fertilization (Fig. 2B, Supplemental Fig. 2G), are morphologically distinct from col1a1a-expressing fibroblasts and were not considered in further analyses. The simultaneous expression of both col1a1a and vimentin, location at the growing fin edge, and cellular morphology led us to speculate that the intricately shaped, double positive cells toward the fin exterior may represent a developmentally matured population of fibroblasts.

### Burn wounding triggers vimentin expression and morphological change similar to changes that occur during fibroblast development

To investigate the response of fibroblasts to burn wound healing, we subjected col1a1a- and vimentin-double positive larvae to burn injury as previously described [18]. Immediately following injury, we observed rounding and eventual loss of fibroblasts in burned tissue (Fig. 3A). While there were approximately 60 col1a1a- and 35-double positive fibroblasts in the region used for burning prior to injury, this number effectively fell to zero by 60 minutes post-burn (mpb) (Figure 3B). To understand the response to acute tissue damage in greater detail, fibroblasts were assessed during the remodeling phase of burn wound repair, starting 24 hours post-burn (hpb) (Fig. 3C). We initially expected the number of fibroblasts in burned tissue to increase over time, corresponding to a greater capacity for ECM remodeling. However, we observed that the total number of fibroblasts already present at 24 hpb did not continue to change over time, with roughly 90 cells present in the burned area from 24 through 72 hpb (Fig. 3D). Given this, we instead hypothesized that the maturation of fibroblasts, indicated by expression of vimentin, may be required for ECM remodeling and burn wound healing. Quantification of double-positive cells over time supported this hypothesis, with the number of fibroblasts expressing vimentin increasing over time (Fig. 3E). At 24 hpb, an average of only 4.3 ± 0.7 cells were double-positive. This more than doubled to 11.4 ± 1.1 cells at 48 hpb, before nearly doubling again to 20.6 ± 1.3 cells at 72 hpb. Importantly, the increasing number of cells expressing vimentin after burn was mirrored by morphological changes reminiscent of mesenchymal cell development described above. As vimentin expression increased over time, we noted that double positive cells had a significantly greater perimeter and reduced circularity, consistent with a more mature phenotype (Fig. 3F, G).

**Figure 3:**
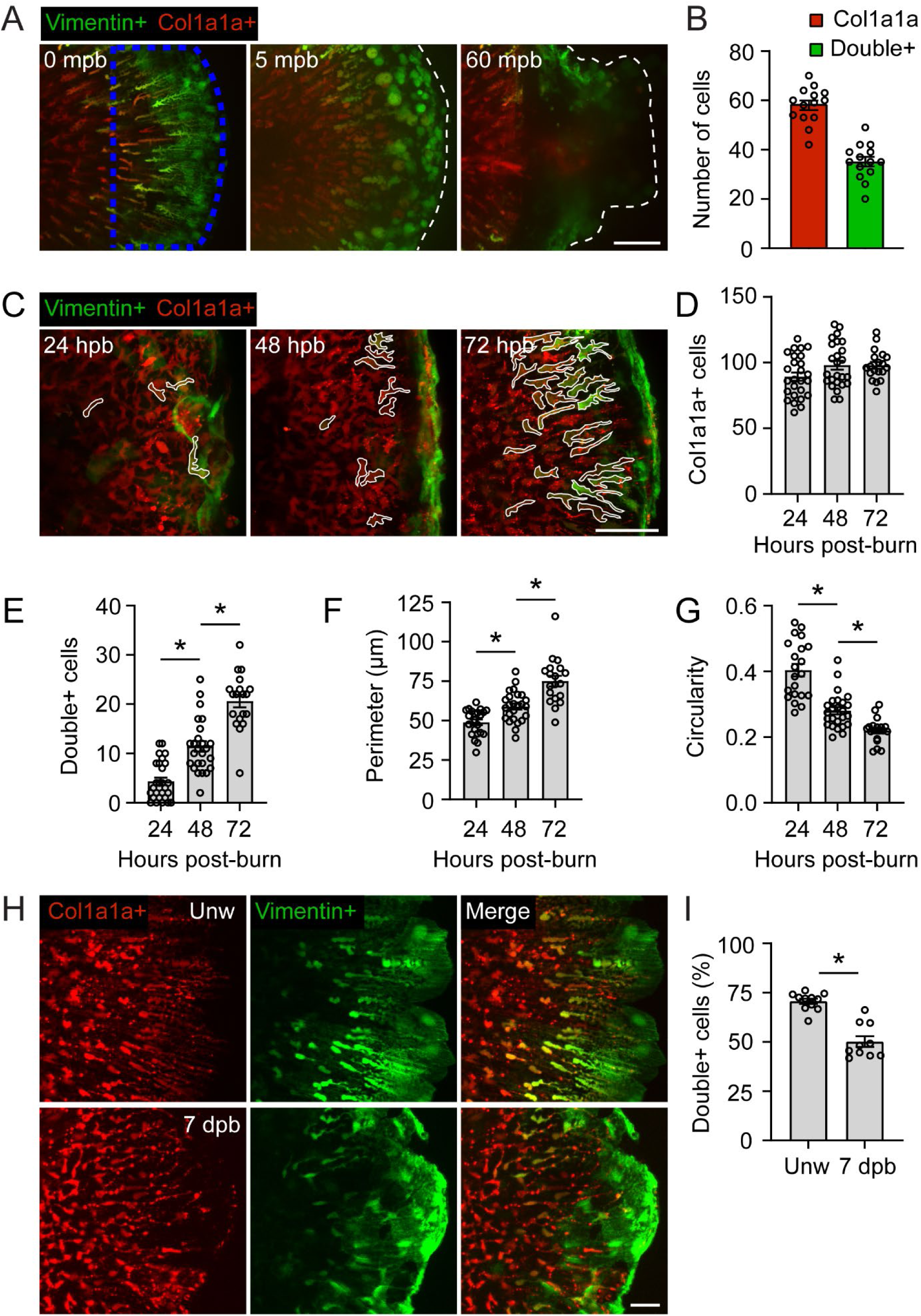
Fibroblasts undergo progressive morphological change and increased vimentin-expression following burn injury. (A) Time series showing the fibroblast response to burn injury (blue dotted region) in a *Tg(Col1a1a:zmCherry)xTg(−2vim:eGFP)* double positive larva over 60 minutes post-burn (mpb). Dotted white line indicates tail fin boundary. Scale bar = 50 µm. (B) Quantification of the total number of col1a1a-only and col1a1a and vimentin double positive cells (double+) in an unwounded fin 3 days post-fertilization (dpf). Region used for quantification corresponds to burned region used in future analyses. (C) Images showing *Tg(Col1a1a:zmCherry)xTg(−2vim:eGFP)* double-positive larvae, outlined in white, at the indicated number of hours post-burn (hpb). Scale bar = 20 µm. (D) Quantification of the total number of col1a1a-positive cells in the burned region at the indicated time. (E) Quantification of the total number of col1a1a and vimentin double positive cells in the burned region at the indicated time. (F) Quantification of the perimeter of col1a1a and vimentin double positive cells. (G) Quantification of the circularity of col1a1a and vimentin double positive cells. (H) Images showing *Tg(Col1a1a:zmCherry)xTg(−2vim:eGFP)* double-positive larvae either unwounded (10 days post-fertilization) or 7 days post-burn (dpb). Scale bar = 20 µm. (I) Quantification of the percentage of col1a1a and vimentin double positive cells in larvae from H. N=14 larvae (B). N ≥ 19 larvae for each time point (D-G). N ≥ 10 larvae for each condition (I). * indicates p<0.05 by One-way ANOVA (E-G) or Mann-Whitney test (I).

Despite the continued increase in vimentin expression over time, we only observed approximately 20 double positive fibroblasts at 72 hpb, corresponding to 21% double positive fibroblasts in the burned region – substantially lower than the 60% of fibroblasts that were double positive prior to burning (Fig. 3B). Burn injuries are known to heal slowly, and in zebrafish there is a significant delay in both inflammation resolution and neuronal regeneration relative to mechanically induced injuries [16, 17]. We hypothesized that fibroblast maturation may experience a similar delay. Indeed, we found that even 7 days post-burn (dpb), the percentage of double positive fibroblasts had still not returned to the level observed in age-matched (10 dpf) unwounded larvae (Fig. 3H, I).

### Vimentin expression is required for fibroblast maturation and burn wound healing

Vimentin is required for normal wound healing [24, 28, 29]. Therefore, we hypothesized that reduced vimentin expression may alter fibroblast behavior and impact burn wound healing. To test this, we decreased vimentin protein level using a previously validated morpholino oligonucleotide (MO) injection [20]. We first ensured that reduction in total vimentin would not alter fibroblast number or gross tail fin development. Compared to controls, vimentin MO injected fish showed no change in the total number of col1a1a-positive, or double positive fibroblasts (Fig. 4A, B), supporting our previous finding that vimentin expression in mesenchymal cells is not an absolute requirement for development [20]. Similarly, there was no difference in tail fin size at 3 days post-fertilization (Fig. 4C).

**Figure 4:**
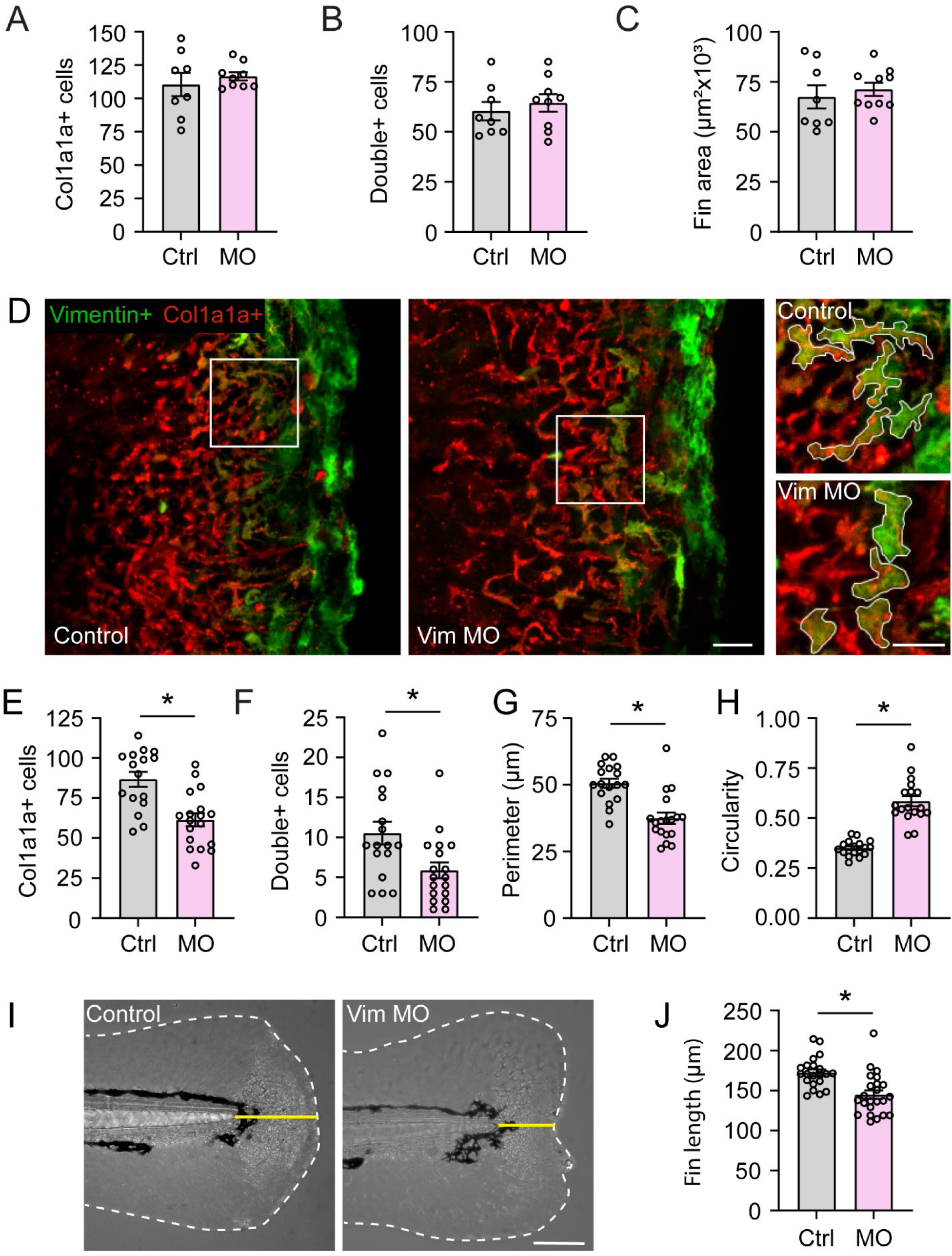
Vimentin-dependent morphological change is required for fibroblast response to burn injury. (A-C) Quantification of the total number of col1a1a-positive cells (A), col1a1a and vimentin double positive cells (B), and tail fin area (C) in unwounded control or vimentin MO injected larvae 3 days post-fertilization. N ≥ 8 larvae for each condition. (D) Images showing control and vimentin morpholino injected *Tg(Col1a1a:zmCherry)xTg(−2vim:eGFP)* double-positive larvae 72 hpb. White box indicates regions shown in inset at right. Double positive mesenchymal fibroblasts are outlined. Scale bar = 20 µm (left) and 10 µm (insets). (E) Quantification of the total number of col1a1a-positive cells in the burned region 72 hpb. (F) Quantification of the number of col1a1a and vimentin double positive cells in the burned region 72 hpb. (G) Quantification of the perimeter of col1a1a and vimentin double positive cells 72 hpb. (H) Quantification of the circularity of col1a1a and vimentin double positive cells 72 hpb. (I) Images showing tailfin morphology of control and vimentin morpholino (Vim MO) injected larvae 72 hpb. Yellow line indicates distance from notochord tip to wound edge used to quantify wound healing. Scale bar = 100 µm. (J) Quantification of wound healing as shown in J. N ≥ 18 larvae for each treatment (E-H). N ≥ 21 larvae for each treatment (J). * indicates p<0.05 by independent t-test.

Larvae were next subjected to burn wound injury with fibroblast maturation assessed at 72 hpb (Fig. 4D). While control larvae had a similar number of col1a1a-positive cells as the uninjected larvae identified above (86.6 ± 4.7 cells), this number was significantly reduced to 61.3 ± 4.1 cells in vimentin MO injected larvae, along with a concomitant decrease in the number of double positive fibroblasts (Fig. 4E, F). Further, in cells that turned on vimentin gene expression, but were unable to produce vimentin protein due to MO injection, we noted a significantly reduced perimeter and increased circularity, reminiscent of a less mature phenotype (Fig. 4G, H). Finally, we tested whether vimentin-dependent fibroblast maturation is required for burn wound healing. For this, we quantified tailfin regrowth 72 hpb and found a 15% decrease in tailfin length in vimentin MO injected fish, representing a significant defect in burn wound healing in the absence of vimentin (Fig. 4I, J). Together, these data indicate that fibroblasts turn on vimentin gene expression in response to tissue injury, mirroring the fibroblast maturation process observed in early development, and this vimentin expression is required for efficient burn wound healing.

### Prolonged neutrophil inflammation inhibits fibroblast maturation in burned tissue

To determine how fibroblasts are affected by the surrounding burn wound environment, we first returned to the RNA-sequencing data. Re-analysis of this data revealed persistent myeloid cell signatures in the burn wounded tissue coincident with increased expression of ECM-related genes (Figure 1). Excessive collagen deposition impairs wound healing and leads to formation of scar tissue, indicating a potentially deleterious overlap of inflammatory and remodeling phases of wound healing. This led us to hypothesize that delayed resolution of inflammation may modulate fibroblast behavior in burned tissue.

To begin testing the link between persistent inflammation and fibroblast function, we treated *Tg(−2vim:eGFP)* larvae with isotonic medium, which significantly limits both neutrophil and macrophage recruitment to burned tissue [18]. Quantification of fibroblast morphology 72 hpb revealed that while total vimentin-positive area was unchanged, fibroblasts labeled by vimentin had a larger perimeter coupled with reduced circularity, indicating a more mature phenotype in larvae treated with isotonic solution (Supplemental Fig. 3A-D). Therefore, it is possible that the restoration of fibroblasts in the isotonic solution treated larvae was at least in part due to reduced immune cell infiltration. Along these lines, we first assessed the potential role of macrophages in modifying fibroblast maturation. Crossing *Tg(−2vim:eGFP)* larvae with the macrophage reporter *Tg(Mpeg-mCherry)* enabled real-time imaging of macrophage-fibroblast interactions after burn injury. While we observed frequent direct physical interactions between these cells 24 hpb, these events were rare by 72 hpb (Supplemental Fig. 4A). While it remains likely that macrophages play a role in regulating fibroblast activity and subsequently ECM remodeling, we instead sought to continue our investigation of neutrophils due to the RNA-seq data suggesting an unexpected persistence of neutrophil related genes at 72 hpb.

To test the role of neutrophils in regulating fibroblast behavior, we utilized *Tg(Mpx:Rac2WT-mCherry)* and *Tg(Mpx:Rac2D57N-mCherry)* larvae [42]. Rac2 is a GTPase that controls neutrophil migration, and neutrophils carrying the D57N mutation are unable to effectively migrate, preventing their recruitment to sites of tissue damage or infection [42, 43]. Burn wounding of Rac2D57N larvae, as well as their control – Rac2WT – counterparts revealed no difference in vimentin-positive cell morphology 24 hpb (Fig. 5A), indicating no early role for neutrophils in modulating fibroblast activity in the inflammatory phase of healing. To determine whether the presence of neutrophils regulates fibroblast behavior during ECM remodeling, we extended our investigation to 72 hpb. Time lapse imaging revealed that neutrophils continue to interact with fibroblasts in the burned region at later time points (Fig. 5B). Notably, these direct interactions often occurred for minutes at a time, with neutrophils shuttling back-and-forth between different fibroblasts. Unlike 24 hpb, preventing neutrophil recruitment to burned tissue affected vimentin-positive cell morphology at 72 hpb (Fig. 5C). As in isotonic medium treated larvae, there was no difference in the total amount of vimentin expressed in the wound area; however, vimentin expressing fibroblasts from Rac2D57N larvae had significantly greater perimeter and reduced circularity (Fig. 5D-F). These findings indicate that neutrophils influence fibroblast function in the burn wound microenvironment, leading us to hypothesize that neutrophil recruitment impacts ECM remodeling in response to burn wound injury.

**Figure 5:**
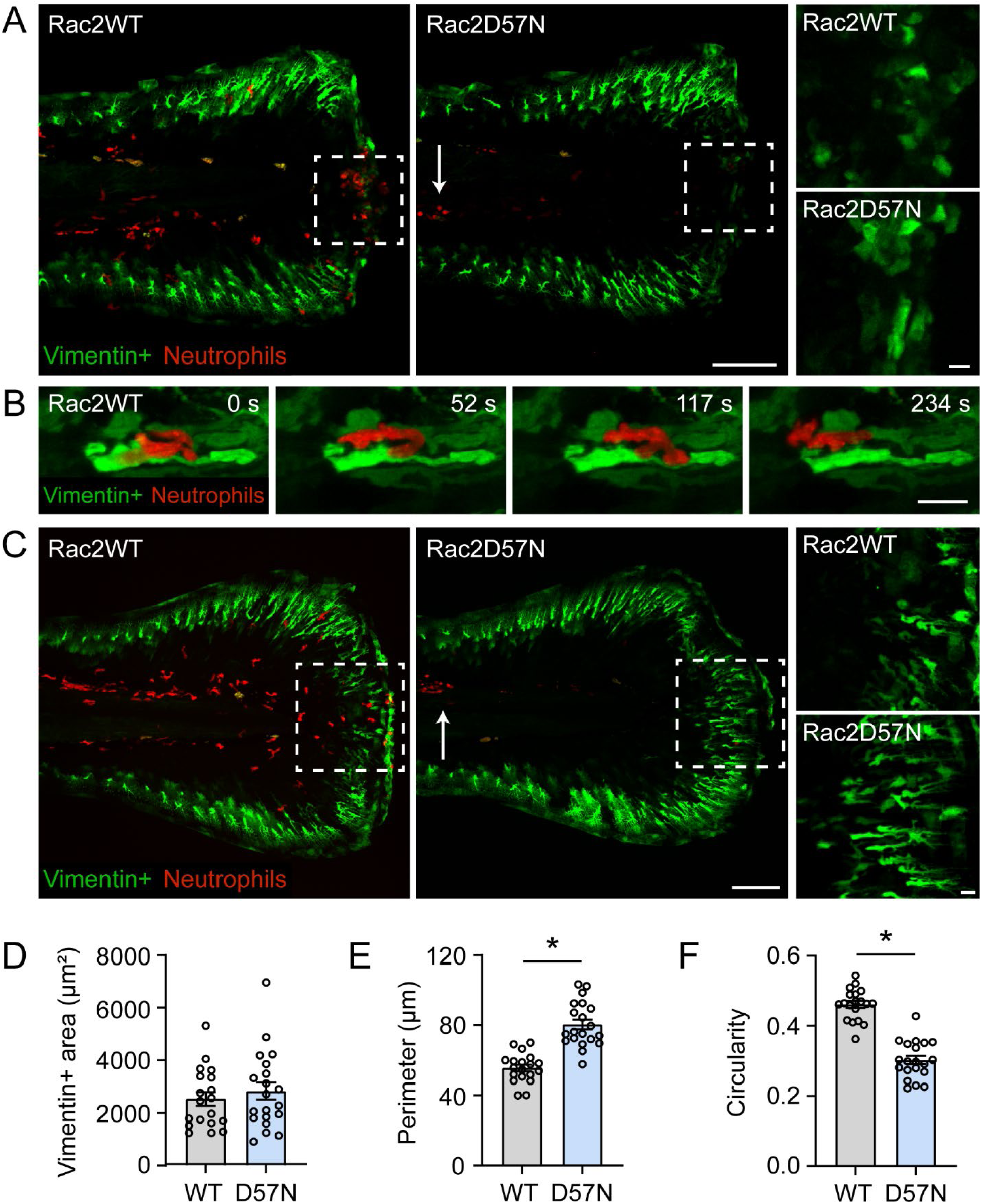
Neutrophils inhibit fibroblast morphological change in burned tissue. (A) Images showing *Tg(−2vim:eGFP)* larvae expressing neutrophil-specific WT *Tg(Mpx:Rac2WT-mCherry)* or D57N mutant *Tg(Mpx:Rac2D57N-mCherry)* Rac2 24 hours post-burn. White box indicates region of inset shown at right. Scale bar = 100 µm (left) and 10 µm (insets). (B) Time series showing a characteristic interaction between a neutrophil and vimentin-positive fibroblasts in *Tg(−2vim:eGFP)*x*Tg(Mpx:Rac2WT-mCherry)* larvae 72 hours post-burn. Scale bar = 10 µm. (C) Images showing *Tg(−2vim:eGFP)* larvae expressing neutrophil-specific WT *Tg(Mpx:Rac2WT-mCherry)* or D57N mutant *Tg(Mpx:Rac2D57N-mCherry)* Rac2 72 hours post-burn. White box indicates region of inset shown at right. Scale bar = 100 µm (left) and 10 µm (insets). (D) Quantification of total vimentin-positive cell area 72 hours post-burn (hpb) in larvae as in C. (E) Quantification of vimentin-positive cell perimeter 72 hpb. (F) Quantification of vimentin-positive cell circularity 72 hpb. N ≥ 20 larvae for each genotype (D-F). * indicates p<0.05 by independent t-test.

### Neutrophil-fibroblast interactions inhibit collagen remodeling after burn injury

We previously showed that burn injury results in an immediate local loss of collagen fibers [20] (Fig. 6A, B). Due to the central role of fibroblasts in the deposition of new collagen following injury, we next used second harmonic generation (SHG) imaging to visualize endogenous collagen in a label-free manner [44]. To test the role of neutrophils in modulating fibroblast function, we assessed collagen fiber regrowth in burn wounded Rac2 WT and D57N larvae using SHG (Fig. 6C). Collagen organization in unwounded fins is characterized by radial outgrowth with fibers growing parallel to each other. Following burn, we found that Rac2D57N larvae had significantly more collagen regrowth 72 hpb and that collagen organization was reminiscent of the healthy, unwounded tail fin (Fig. 6D), supporting our hypothesis that neutrophils inhibit fibroblast function. Importantly, we noted disorganized collagen regrowth in burned fish, indicating a significant delay in ECM remodeling. Therefore, we next asked whether the absence of neutrophils affected collagen organization. For this, blinded analysis of SHG images was performed with images ranked into categories of (1) no defect (fibers growing radially as in unwounded fins), (2) mild defect (disorganized fiber regrowth is restricted to small patches taking up a minority of the burned region), or (3) severe defect (disorganized fiber regrowth is widespread and takes up a majority of the burned region) (Fig. 6E, F). This revealed a significant difference between Rac2 WT and D57N larvae with significantly more Rac2 D57N larvae exhibiting no observable defect in collagen remodeling. In fact, 92.1% (35 out of 38 total rankings) of all images ranked in category 1 were Rac2 D57N larvae. Interestingly, while images receiving a score of 3 made up a minority of total rankings, only 18.5%, there was no meaningful difference in the distribution of WT and D57N. Automated analysis of collagen fibers supported the blinded qualitative ranking with Rac2 WT larvae showing significantly larger deviation angles in collagen fiber alignment in burned tissue (Fig. 6 G, H) [45, 46]. Thus, prolonged neutrophil infiltration of burned tissue contributes to defects in collagen ECM remodeling, likely by altering collagen-expressing fibroblast function.

**Figure 6:**
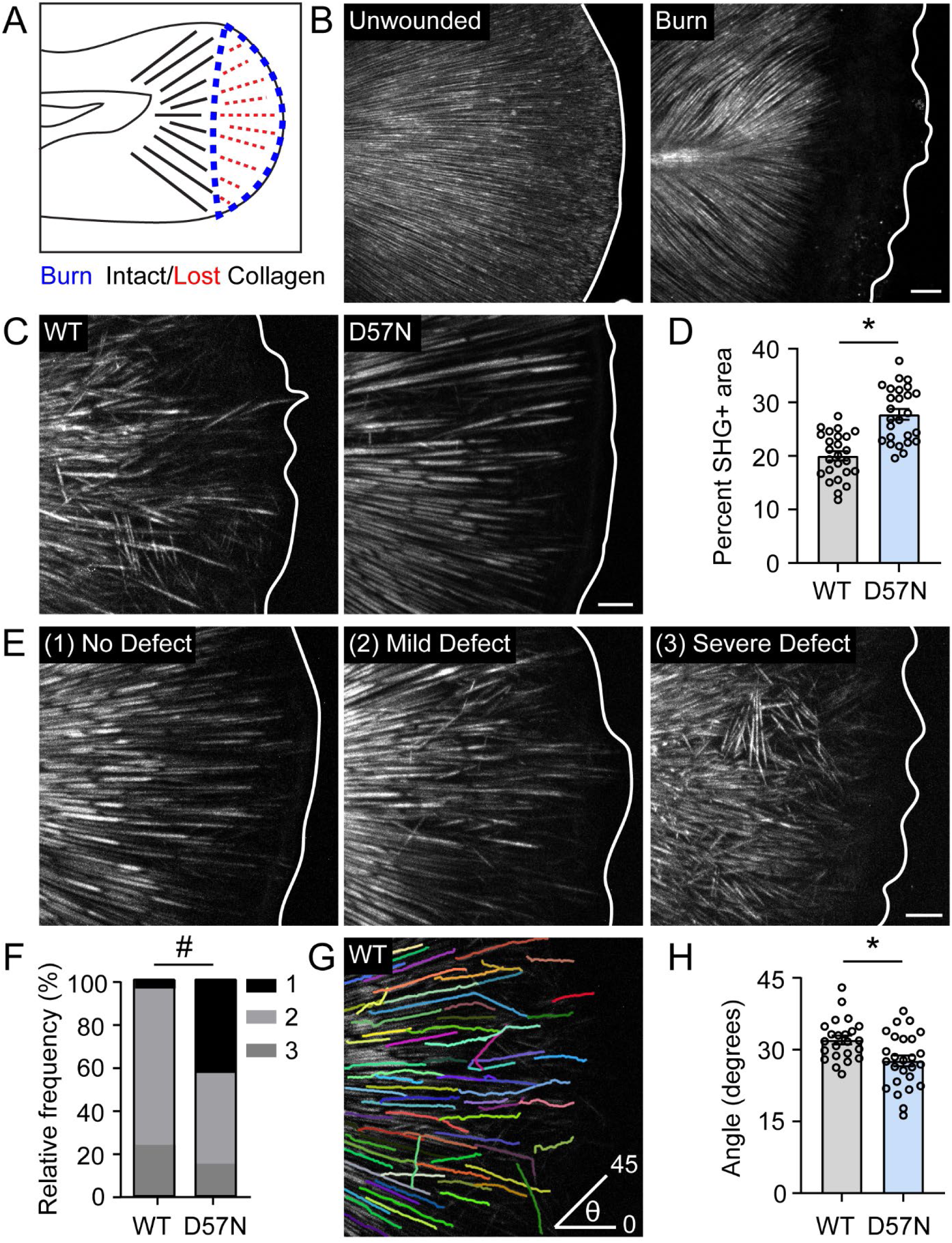
Neutrophils inhibit collagen extracellular matrix remodeling during burn wound healing. (A) Schematic illustrating the consequence of burn wound injury on collagen fibers in the larval zebrafish tailfin. (B) Second harmonic generation (SHG) images of unwounded or burn wounded larvae 5 minutes post-burn (mpb). (C) SHG images showing *Tg(Mpx:Rac2WT-mCherry)* and *Tg(Mpx:Rac2D57N-mCherry)* 72 hours post-burn. (D) Quantification of the total SHG-positive area 72 hours post-burn. (E) Representative SHG images showing examples of collagen remodeling considered to result in No Defect, Mild Defect, or Severe Defect. (F) Quantification of blinded ranking of representative SHG images into categories defined in E. (G) Image depicting the method used to quantify collagen fiber angle in burned tissue. (H) Quantification of collagen fiber angle 72 hours post-burn. In all cases, solid white line denotes tailfin boundary. N ≥ 25 larvae for each genotype (D, H). * indicates p<0.05 by independent t-test. ^#^ indicates p<0.05 by Mann-Whitney test. In all cases, scale bar=20 µm.

## Discussion

Fibroblasts are critical for tissue development and repair. However, the behavior of fibroblasts in living tissues remains unclear, in part, because of the lack of tools to image fibroblasts in real time. Here we generated new fluorescent reporter zebrafish lines to investigate fibroblast behavior during development and in response to tissue injury. We find that expression of vimentin in col1a1a positive fibroblasts correlates with morphological changes associated with a more developmentally matured phenotype. In response to burn injury, fibroblasts take on an elongated morphology and progressively increase expression of vimentin over time, mirroring their maturation during larval development. Prolonged and excessive neutrophil infiltration inhibits fibroblast maturation and collagen remodeling, resulting in delayed ECM remodeling that can be rescued by preventing neutrophil infiltration of burned tissue.

Dissecting the role of fibroblasts has proven challenging due to the functional heterogeneity they display within tissues. Distinct subpopulations of fibroblasts contribute to tissue development and wound healing in mammalian skin, and the lack of spatial segregation of each subpopulation makes interrogating their individual functions challenging [47, 48]. Larval zebrafish provide a unique opportunity to expand our understanding of how fibroblast identity and spatial localization within tissue impacts function during development and wound healing in vivo. For example, studies of fibroblasts during zebrafish development have identified the requirement for early mesoderm-derived fibroblasts in fin morphogenesis. Early mesoderm cells populate the developing fin fold starting on the first day post-fertilization [9], and must take on progressively more elongated morphologies over time [40], suggesting that precise migratory dynamics are coupled with fibroblast maturation during fin morphogenesis. Through real-time imaging of larval development using dual fibroblast reporter zebrafish, our work adds to this understanding of dynamic fibroblast behavior during development *in vivo*. We identify that fibroblasts only begin to take on morphologies associated with an activated phenotype approximately 35 hours post-fertilization when vimentin expression is turned on in the outermost fibroblasts of the developing tail fin. Curiously, vimentin is never expressed in fibroblasts closest to the midline, with vimentin-positive fibroblasts continually expanding outward as the fin grows. This suggests that during development, fibroblasts are only actively remodeling ECM at the outer edge of the tissue as the tailfin grows. Additional work is required to understand the dynamics of these vimentin positive cells over time. For example, it is possible that expression of vimentin during development only occurs once, with newly activated fibroblasts continually expanding outward as the fin grows. However, an unexplored alternative hypothesis is that activated fibroblasts lose vimentin expression, allowing for a new set of activated fibroblasts to take over as the tissue expands outward. Interestingly, vimentin has been shown to label all mesenchymal-derived fibroblasts in adult human skin [48]. This indicates potential species-specific differences in the regulation of fibroblast identity, but also opens the possibility that vimentin plays a role in regulating fibroblast development, which may be explored further using the zebrafish model generated here.

In response to tissue damage, it is well-established that fibroblasts take on an activated phenotype (often called myofibroblasts in mammals) [2, 49]. However, chronically activated fibroblasts are associated with fibrosis. More recently, evidence suggests that fibroblast activation is transient [2, 41], but the regulation of this transient state is not well-understood. Here we leverage intravital imaging, using vimentin expression and fibroblast morphology, to pinpoint the spatial and temporal kinetics of fibroblast activity following tissue injury. Of note, we find that fibroblasts must repopulate the burned region before they become fully activated (Figure 3A, C). This timing, beginning 24 hpb, corresponds to the onset of the remodeling phase of burn wound healing after the accumulation of tissue damage and initial immune cell recruitment has abated [16, 18]. Similar to our observations from developing larvae, fibroblast maturation occurs preferentially toward the outer bound of the wound edge. Collectively, these observations suggest that immature fibroblasts either proliferate or migrate to the site of injury prior to undergoing activation. Interestingly, we found that fibroblast expression of vimentin was not restored relative to age-matched unwounded larvae by 7 days post-burn (Fig. 3H), highlighting the significant delay in restoring the skin to homeostasis following burn injury, despite the overall tissue appearing visually normal. Our findings also imply that mechanical or biochemical signals that regulate fibroblast activation are present in the surrounding wound microenvironment, similar to the finding that mechanosensitive *Engrailed-1* expression that contributes to fibrosis is only present in wound-exposed fibroblasts [50]. In addition to environmental signaling, the fibrogenic potential of fibroblasts has also been shown to depend on their developmental origin. In mammals, fibroblasts of the neural crest that populate facial skin showed increased expression of Robo2 and subsequent downstream effectors, decreasing their scarring potential relative to mesoderm derived fibroblasts [51]. Using real-time imaging to interrogate the mechanisms that regulate vimentin expression and fibroblast identity in zebrafish during development may provide an opportunity to dissect the cues that regulate fibroblast function during tissue repair, a time when environmental signaling is significantly more complex.

Scar formation and subsequent loss of tissue function represents a growing problem affecting burn wound patients; however, the underlying mechanisms resulting in scar formation are not well understood. While fibroblasts are the cells primarily responsible for ECM deposition, both fibroblasts and immune cells, such as neutrophils, secrete matrix metalloproteinases (MMP) that degrade and remodel ECM [1, 52, 53]. While our data implicates infiltrating neutrophils in modulating fibroblast activity and function, the mechanism by which neutrophils alter ECM remodeling are not clear. Our data suggests that early fibroblast activation is independent of the neutrophil response, with vimentin expression 24 hpb being unchanged in the absence of infiltrating neutrophils (Figure 5A). This indicates that the initial influx of neutrophils is not detrimental to fibroblast function, but prolonged neutrophil inflammation impairs fibroblast activity. In the current study we chose to focus on neutrophils due to the unexpected persistence of neutrophils in the burn wound environment. However, burn wounding larval zebrafish also generates a robust macrophage response ([16], Supplemental Fig. 4), and macrophage-fibroblast interactions have been well characterized in mammals [54]. In response to tissue damage, macrophages are attracted to the site of damage where they play a role in negatively regulating infiltrating neutrophils, and in this capacity, they also directly contact fibroblasts [55]. Highlighting the importance of direct contact, it has been shown that macrophage contact can activate fibroblasts in a mechanism dependent on the mechanosensitive channel Piezo1, contributing to fibrosis [56]. Interestingly, we observed direct contact between macrophages and vimentin-expressing fibroblasts 24 hpb; however, these interactions were largely absent by 72 hpb, with macrophages receding from the burned region (Supplemental Fig. 4). This may indicate the presence of an endogenous regulatory network between neutrophils, macrophages, and fibroblasts that controls the duration of immune cell-fibroblast contacts in an effort to restore homeostasis.

The RNA-sequencing data reported here provides a means of identifying potential pathways by which neutrophils may indirectly regulate fibroblast activity. One possibility is through increased MMP expression. MMPs were broadly increased throughout burn wound healing (Supplementary Table 1), including MMP9, which is highly expressed in neutrophils. Further, MMP9 has been shown to be upregulated in burn patients [57], with neutrophil-derived MMP9 proposed to contribute to tissue damage [58]. Therefore, it is possible that prolonged neutrophil accumulation in burned tissue results in excessive local MMP secretion and collagen degradation. In this way, chronic neutrophil inflammation may result in scar formation due to an imbalance in ongoing ECM deposition by fibroblasts with simultaneous degradation by neutrophils. However, future work toward therapies targeting neutrophil infiltration in burned tissue must focus on alleviating the prolonged nature of inflammation without disrupting the ability of neutrophils to be recruited to burned tissue at all. Preventing neutrophil infiltration entirely may prove deleterious, not only due to the positive role neutrophils play in regulating fibroblast collagen production [35], but also due to the critical role of neutrophils in fighting infection, which is the primary driver of burn patient mortality [59–61]. While our RNA-sequencing may provide unique insights to burn pathology as described above, it is important to note that bulk RNA-sequencing as performed here is unable to distinguish cell type-specific expression of genes and thus must be interpreted with caution. Indeed, even expression of vimentin, a characteristic marker of fibroblast identity, is notably expressed in cells of the outer fin edge (Fig. 2E). Thus, we are unable to completely rule out a contribution of these cells in ECM remodeling during wound healing or development. Future work generating reporter zebrafish must focus on identifying individual fibroblast subpopulations with greater specificity.

Our current work resulted in new fluorescent reporter zebrafish to investigate fibroblast behavior during development and tissue repair. In doing so, we identified that fibroblast behavior in burned tissue mirrors the normal maturation of fibroblasts during tailfin development but is augmented by excessive inflammation. This observation suggests wound-associated environmental factors alter fibroblast behavior and promote scar formation. Future studies of fibroblast behavior during development may therefore lay a foundation for understanding the optimal regulation of fibroblast activation and provide a basis for understanding how fibroblast activation becomes dysregulated in response to tissue injury.

## Materials and Methods

### Animal Handling and Ethics

Zebrafish studies were carried out in accordance with the recommendations from the Guide for the Care and Use of Laboratory Animals. All zebrafish experiments performed were approved by the University of Wisconsin – Madison Research Animals Resource Center under the Protocol M005405-R02. Adult fish were maintained on a 14-hour light, 10-hour dark cycle. Following breeding, embryos were transferred to E3 medium (5 mM NaCl, 0.17 mM KCl, 0.44 mM CaCl_2_, 0.33 mM MgSO_4_, 0.025 mM NaOH, and 0.0003% methylene blue) and maintained at 28.5°C until 3 days post-fertilization. For all experiments, 3 days post-fertilization larval zebrafish were anesthetized in 0.2 mg/mL tricaine (ethyl 3-aminobnzoate) in E3 medium prior to use. Treatment with isotonic medium (E3 supplemented with NaCl to a final concentration of 135 mM) was performed as previously described [17]. Larvae were exposed to isotonic medium immediately prior to burn injury and washed into fresh E3 24 hours post-burn. When necessary, larvae were screened for positive fluorescence using a Zeiss Zoomscope EMS3/SyCoP3 with a Plan-NeoFluar Z objective. All transgenic lines including *Tg(Col1a1a:zmCherry)*, *Tg(Col1a1a:acGFP) Tg(Col1a1b:acGFP)*, *Tg(Col1a2:acGFP)*, *Tg(−2vim:eGFP)* [20], *TgBac(Lamc1:Lamc1-sfGFP)* [62], *Tg(Mpx:Rac2WT-mCherry)* [42], and *Tg(Mpx:Rac2D57N-mCherry)* [42] were maintained on the AB background strain.

### Generation of collagen reporter fish lines

To generate transgenic reporter zebrafish for collagen 1 expression, the upstream region of the putative translational start site of the zebrafish collagen 1a1a (3 kb upstream), collagen 1a1b (1.5 kb upstream), and collagen 1a2 (2.5 kb upstream) were PCR amplified using the following primers: Col1a1a – F: TAAGTGCTGTATAGTGGTTCGC R: CTTTGAGGCGAGGGAAGTTC Col1a1b – F: CGTAAAGATGTGAGAAAGGAGG R: GACATGTAGACTCTTTGAGGC Col1a2 – F: TGCTTGATAGCAAAGTTCTACACA R: GTCTAAACCGACATGCAGAC

The PCR amplified products were cloned into Tol2 expression vectors containing *acGFP* or *zmCherry* for fluorescence labeling and used for genomic integration [63]. F_0_ larvae were screened for positive fluorescence using a Zeiss Zoomscope EMS3/SyCoP3 and crossed for two generations to obtain stable reporter lines.

### Burn wounding of larval zebrafish

To perform burn injury, zebrafish larvae were transferred to a 60 mm tissue-culture treated dish containing 0.2 mg/mL tricaine in E3 medium. A fine tip cautery pen (Geiger Medical Instruments) was used to burn the caudal fin of larval zebrafish. Burn wounds were applied until the wounded region bordered but did not touch the posterior notochord boundary. In all cases, larvae were washed to remove tricaine immediately following injury until ready for use. To quantify burn wound healing, the shortest straight-line distance from the posterior tip of the notochord to the burn wound edge was measured.

### Tissue collection and RNA-sequencing

Tail fins from 50 unwounded or burn wounded larvae were pooled and collected in ice cold PBS for each time point. Three biological replicates, representing unique breeding clutches, were collected for each time point. RNA was extracted from pooled fins using TRIzol reagent and RNAqueous Micro Kit (Invitrogen). Extracted RNA was submitted to GENEWIZ for library preparation and sequencing. Pooled RNA libraries were sequenced on an Illumina HiSeq to obtain 150 bp paired end reads.

### RNA-seq data processing and differential expression analysis

Raw sequencing data were aligned to zebrafish genome GRCz11 with an improved gene annotation [64] using RSEM [65] with parameters “-p 16 --paired-end --star --estimate-rspd --append-names --output-genome-bam” and STAR version 2.7.11b [66]. Post alignment, gene-level expression estimates were used for differential expression analysis with DEseq2 (v1.38.3) [67]. For visualizing differentially expressed genes on volcano plots, log2 fold changes between conditions were shrunk with *lfcShrink* (type = “ashr”) [68], and p-values were adjusted by the Benjamini-Hochberg method [69].

### Weighted gene co-expression network analysis

Top 25% variance genes were included for a single-block WGCNA analysis, with parameters “corType = “pearson”, power = 12, networkType = “signed”, mergeCutHeight = 0.25, numericLabels = T, minModuleSize = 30, maxBlockSize = 6000” [37]. Power level (power = 12) was determined through soft threshold estimation, while merging height (mergeCutHeight = 0.25) was determined based on adjacency estimation. Genes associated with each module were converted to Entrez gene ID and used as input for Gene Ontology enrichment analysis through Metascape (V3.5.20250701) focusing on GO Biological Processes [38].

### Vimentin morpholino injection

Morpholino oligonucleotides targeting vimentin transcripts (Exon 1 – Intron 1) with the sequence 5’ – GTAATAGTGCCAGAACAGACCTTCTC – 3’ was used as previously published [20]. 3 nl of vimentin morpholino at a concentration of 80 µM was injected into the yolk of an embryo at the one-cell stage. Control injections were performed exactly as morpholino injections with sterile water replacing vimentin morpholino. Larvae were kept at 28.5° until 3 days post-fertilization.

### Microscopy and image analysis

For live imaging, larvae were either mounted in a zWEDGI restraining device allowing for tailfin movement [70, 71] or entirely immersed on a 35 mm glass bottom dish (CellVis) using 1% low-melting point agarose (Sigma-Aldrich). All imaging was performed using a Nikon Eclipse Ti2 microscope equipped with a Crest X-light V3531 spinning disc unit and a Kinetix22 monochrome camera. Imaging was performed using Plan Apochromat 10X/0.45 NA, S Fluor 20X/0.75 NA, S Fluor 40X/1.30 NA, and Plan Apochromat 60X/1.42 NA objectives. All image analysis was performed using ImageJ.

### Quantification of mesenchymal cell morphology

To assess cell morphology, a 150 µm^2^ region of interest at the center of the burn wounded tissue was cropped. Mesenchymal cells were manually outlined using the polygon tool in ImageJ and area, perimeter, and shape descriptors (circularity) measurements were calculated.

### Multiphoton microscopy of second harmonic generation and image analysis

Fixed caudal fin samples from PTU-treated larvae were prepared as previously described [72, 73] by removal from the body with a scalpel blade (Feather #15) then imaged in a 50 mm cover glass (#1.5) bottom dish (MatTek, Ashland MA). The glass bottom depression was covered with a second 18×18 mm #1.5 coverslip to minimize sample movement. Caudal fins were imaged using a custom-built multiphoton microscope [72, 74] at the Laboratory for Optical and Computational Instrumentation using a 20X PlanApo air immersion lens (0.75 NA, WD = 1.0) (Nikon, Melville NY) on a Nikon TE2000 inverted microscope. The multiphoton excitation source Ti:Sapphire laser (Chameleon UltraII, Coherent Inc., Santa Clara, CA) was tuned to 890 nm and the backwards SHG was collected using a 445/20 nm bandpass emission filter (Semrock, Rochester NY). The signal was detected using a H7422P-40 GaAsP Photomultiplier Tube (PMT) (Hamamatsu, Japan). Images were collected using OpenScan software (https://loci.wisc.edu/openscan/). Transmitted-light images were collected with a photodiode-based transmission detector (Bio-Rad, Hercules CA). For each caudal fin, data were collected as z-stacks at two zooms: 0.5x zoom to include the notochord, with 4 µm between sections and one scan per section, and 1x zoom of the fin edge, with 2 µm between sections and 8 scans per section, which were averaged in FIJI [75] for improved image quality. All images were collected at 512 x 512 resolution. Ranked scoring of SHG images was performed by three volunteers. Volunteers were blinded to the genotype of each max-projected SHG image and asked to score the image into a category based on the following criteria: (1) Minimal/No Defect – There are only radially organized collagen fibers or few (1-2) non-radial fibers. (2) Mild Defect – There are mostly radially organized fibers, but interspersed are haystack fibers, either in patches or broadly covering a region of the tailfin. (3) Severe Defect – Much of the burned region contains non-radial, haystack organization of collagen fibers. Scores from each volunteer were pooled and used to generate a relative frequency of each score per genotype. For quantitative analysis of collagen fiber organization, fiber alignment angles were computed using CT-FIRE module of CurveAlign (v5.0 Beta, https://loci.wisc.edu/curvealign/) running on Matlab 2018b. Raw images were imported and fibers identified in batch mode using the curvelet transform [45, 46] with minimum fiber length of 75 pixels (∼27.5 µm) and maximum fiber width of 15 pixels (∼5.5 µm). All angles were measured as the amount of deviation (in degrees) from the image horizontal axis, so that purely horizontal fibers have an angle of 0° and purely vertical lines an angle of 90°.

## Statistical analysis

All statistical analysis and graphing were performed using GraphPad Prism version 10.3.0. The D’Agostino and Pearson normality test was used to determine the appropriate statistical test based on normal distribution of data. For comparison of two means, independent t-tests were used for normally distributed data and Mann-Whitney test was used for non-parametric data. For multiple comparisons, one-way ANOVA with Tukey’s multiple comparison test was used for normally distributed data. For comparison of ordinal data from blinded ranking analysis, the non-parametric Mann-Whitney test was used. Unless otherwise indicated, all data are represented as mean ± SEM (standard error) with p<0.05, indicated by asterisk, used to determine statistical significance. All experiments were performed with at least 3 independent biological replicates defined as either separate breeding clutches of zebrafish or human subjects, with the N-value used for statistical analysis indicated within the relevant figure legend.

## Acknowledgements

The authors would like to acknowledge K99GM138699 to V. Miskolci and R35GM118027-05 awarded to A. Huttenlocher. We would like to thank the members of the Huttenlocher lab for their careful reading of this manuscript and helpful comments.

## Data and Resource Availability

All raw and processed sequencing data generated in this study have been submitted to the NCBI Gene Expression Omnibus (GEO; http://ncbi.nlm.nih.gov/geo/) under accession number GSE315881. The data can be accessed by reviewers with the token: irulsuqmrpsvvob. Bulk RNA-seq analysis codes are available at: https://github.com/k326xh/zebrafishburn. All data, including new fish lines generated as part of this study, will be made available upon request.

