## Supplementary Information for "Fibroblast response to burn injury in larval zebrafish mirrors developmental maturation and is inhibited by infiltrating neutrophils"

Supplemental Figure 1

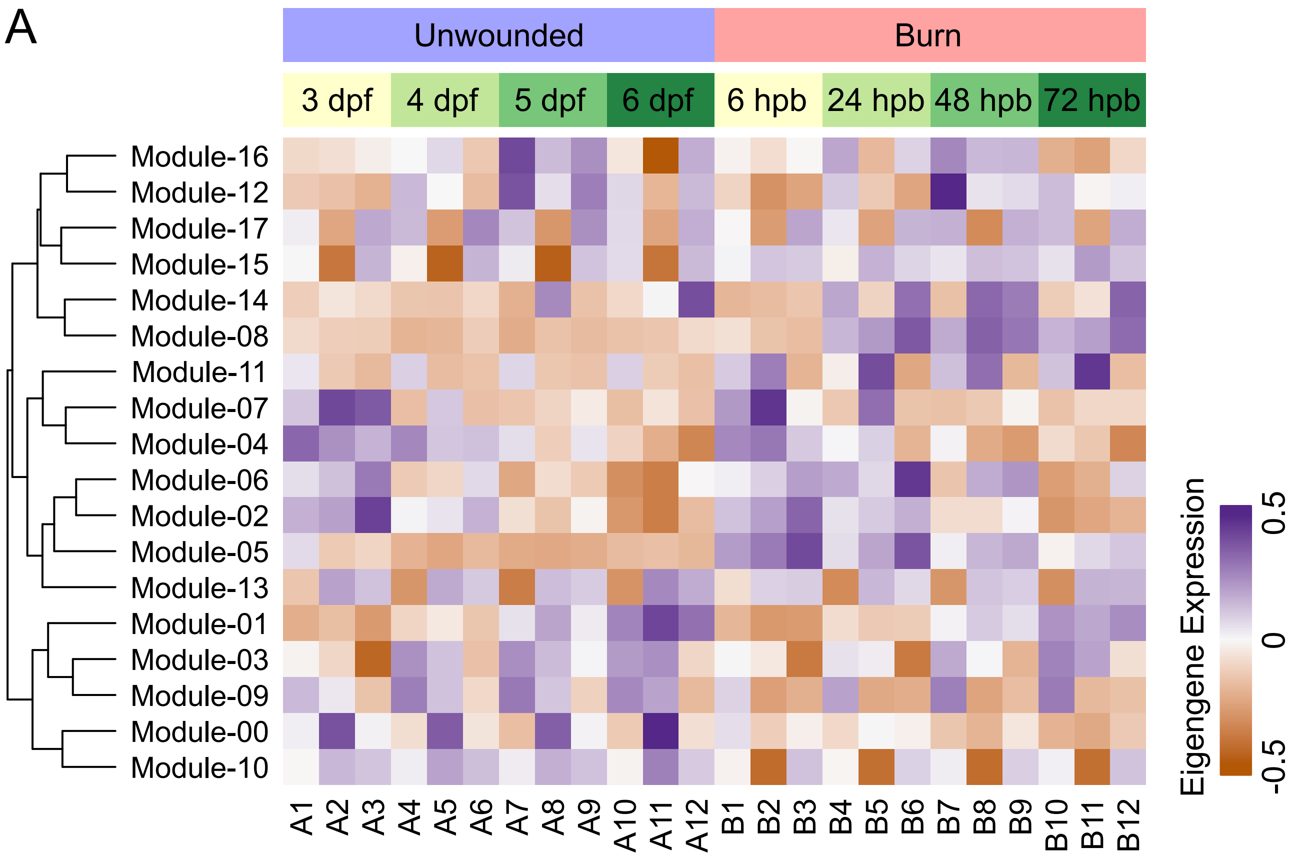

### Supplemental Figure 2

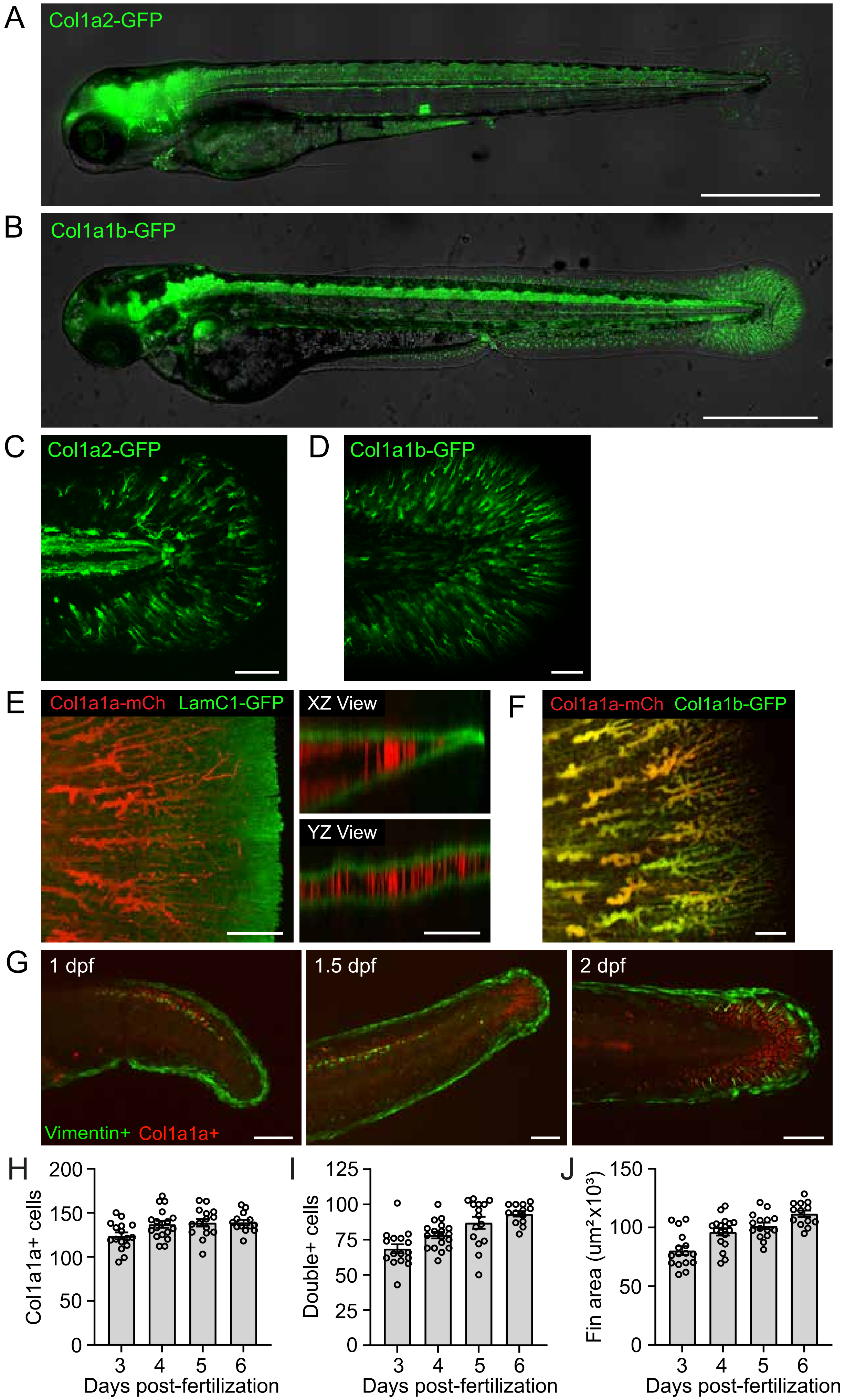

Supplemental Figure 3

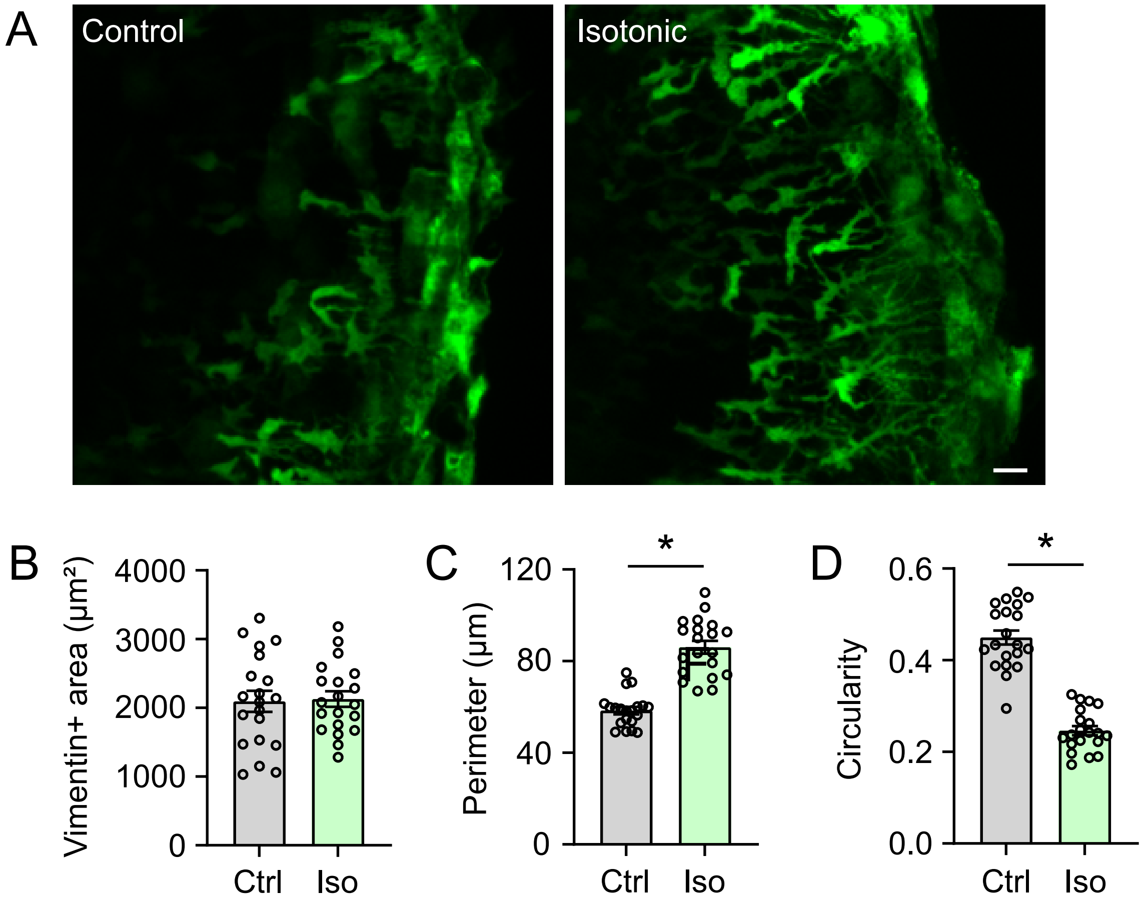

### Supplemental Figure 4

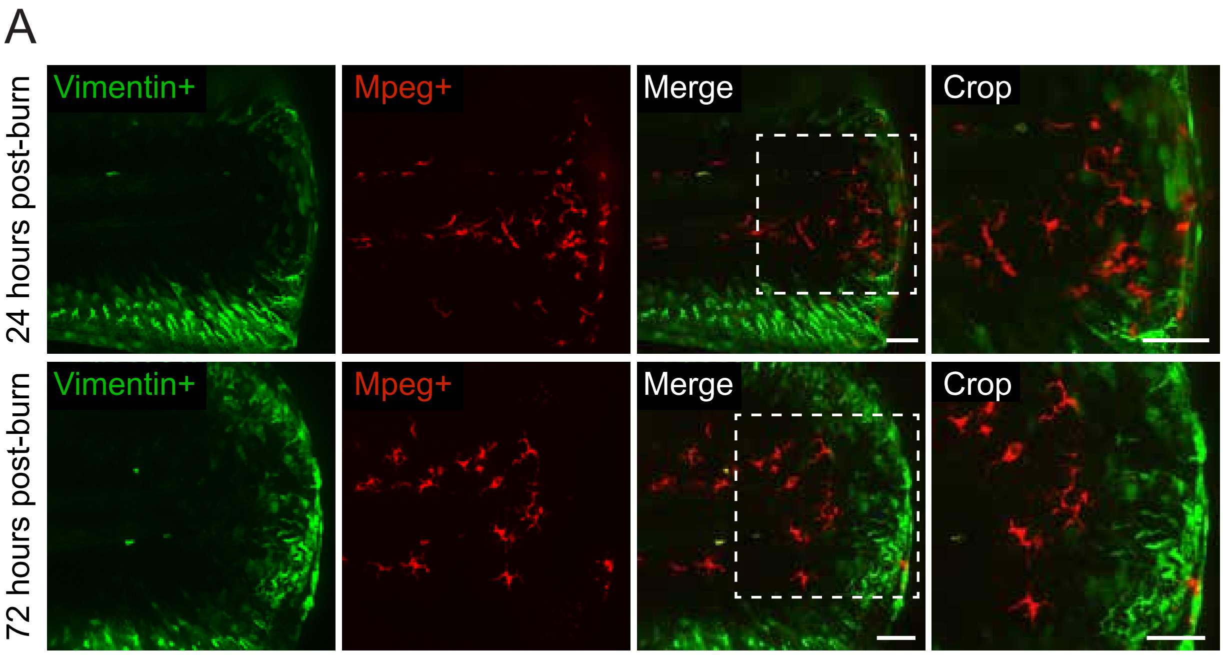

**Supplemental Information**

**Supplemental Figure 1: . Gene modules governing the burn wound healing process.** (A) Heatmap of module eigengene expression across samples. Biological replicates are grouped by wounding condition and time of sample collection.

**Supplemental Figure 2: Collagen I labels a mesenchymal cell population localized to the developing zebrafish dermis.** (A) Image showing *Tg(Col1a2:acGFP)* larvae 3 days post-fertilization. Scale bar = 500 µm. (B) Image showing *Tg(Col1a1b:acGFP)* larvae 3 days post-fertilization. Scale bar = 500 µm. (C) Image showing *Tg(Col1a2:acGFP)* labeled mesenchymal cell population in the tailfin. Scale bar = 100 µm. (D) Image showing *Tg(Col1a1b:acGFP)* labeled mesenchymal cell population in the tailfin. Scale bar = 100 µm. (E) Image showing the tailfin of *Tg(Col1a1a:zmCherry)xTgBac(Lamc1:Lamc1- sfGFP)* larvae. Scale bar = 20 µm. (F) Image showing co-localization of fluorescence signal in mesenchymal fibroblasts in *Tg(Col1a1a:zmCherry)xTg(Col1a1b:acGFP)* larvae. Scale bar = 10 µm. (G) Time series showing developmental origin of *Tg(Col1a1a:zmCherry)xTg(-2vim:eGFP)* labeled mesenchymal cells from 1-2 days post-fertilization. Scale bar = 100 µm. (H-J) Quantification of the number of col1a1a-positive cells (H), col1a1a and vimentin double positive cells (I), and tail fin area (J) in unwounded larvae at the indicated day post-fertilization. N ≥ 13 larvae for each day.

**Supplemental Figure 3: Treatment with isotonic solution promotes fibroblast morphological change in burned tissue.** (A) Images showing *Tg(-2vim:eGFP)* larvae 72 hours post-burn. Larvae were treated with isotonic solution for 24 hours post-burn. Scale bar = 10 µm. (B) Quantification of total vimentin-positive cell area 72 hours post-burn (hpb) in larvae as in A. (C) Quantification of vimentin-positive cell perimeter 72 hpb. (D) Quantification of vimentin-positive cell circularity 72 hpb. N ≥ 20 larvae for each treatment (B-D). * indicates $p<0.05$ by independent t-test.

**Supplemental Figure 4: Direct, physical interaction between macrophages and vimentin-positive fibroblasts is restricted to early burn wound healing.** (A) Images showing *Tg(-2vim:eGFP)xTg(Mpeg-mCherry)* larvae 24- and 72-hours post-burn. Macrophages (Mpeg+) often physically interact with vimentin+ fibroblasts at 24 hours post-burn, but physical interactions were rarely observed by 72 hours post-burn. White dashed box indicates cropped area shown at right. Scale bars = 50 μm.

**Supplemental Table 1: List of differentially expressed genes comparing unwounded and burned tissue.**

**Supplemental Table 2: Functional annotation of WGCNA-calculated gene modules associated with burn wound healing.**

**Supplemental Movie 1: Neutrophil fibroblast interaction in burned tissue.** Neutrophil *Tg(Mpx-Rac2WT:zmCherry)* interacting with two fibroblasts *Tg(-2vim:eGFP)* 72 hours post-burn injury. Neutrophils routinely engage in prolonged physical interaction with wound-responsive fibroblasts. Images were collected at 60x magnification with a 13 second interval. Scale bar = 10 μm. Time is recorded in mm:ss.
